# Oxidative stress induced by tBHP in human normal colon cells by label free Raman spectroscopy and imaging. The protective role of natural antioxidants in the form of β-carotene

**DOI:** 10.1101/2021.02.16.431391

**Authors:** B. Brozek-Pluska, K. Beton

## Abstract

The present study aimed to investigate the protective effect of β-carotene on the oxidative stress injury of human normal colon cell line CCD-18Co triggered by tert-Butyl hydroperoxide (tBHP). XTT examination was used to determine cell viability after β-carotene supplementation and to determine the optimal concentration of antioxidant in spectroscopic studies. Cell biochemistry for CCD-18Co control group, after tBHP adding and for cells in β-carotene - tBHP model was studied by using label-free Raman microspectroscopy. Results for stress treated CCD-18Co human colon normal cells and human colon cancer cells Caco-2 based on vibration features were also compared. Pretreatment with β-carotene alleviated damages in CCD-18Co human normal colon cells induced by tBHP and showed the preventative effect on cells apoptosis. Treatment with β-carotene altered the level of ROS investigated based on intensities of Raman peaks typical for lipids, proteins and nucleic acids. Presented study confirmed the antioxidant, protective role of β-carotene against ROS by using spectroscopic label-free Raman techniques.

## Introduction

Carotenoids, also called tetraterpenoids are yellow, red and orange pigments produced by plants, fungi as well as bacteria. They are responsible for specific and well recognizable color of e.g. pumpkins, carrots, corn, tomatoes, flamingos, salmon and lobsters. There are about 600 plant pigments in the world, but only six of them have a significant impact on human body - α- and β-carotene, β-cryptoxanthin, lutein, lycopene and zeaxanthin. Up to now, around thousand known carotenoids and their derivatives are categorized into two major classes, xanthophylls (including oxygen in the molecular structure) and carotenes (which are purely hydrocarbons). In plants carotenoids are accumulated in the plastids (chromatophores and chromoplasts) and play important role as structural and functional compounds. They are inactive in photosynthesis but play an auxiliary role as light-harvestinging pigments, absorbing light energy for use in photosynthesis; simultaneously they provide photoprotection via non-photochemical quenching minimizing the results of photooxidative stress [1–3].

Mammals obtain carotenoids predominantly through plant foods, carnivorous animals obtain them also from animal fat [4]. In organism carotenoids are absorbed by the intestines into the blood, which transports them to various tissues in the body using lipoproteins. Approximately twenty of fat-soluble carotenoids are found in human blood and tissues [5]. In mammals β-carotene, vitamin E and other carotenoids are commonly perceived as antioxidants, but their biological activities are different and distinguish them from each other [6,7]. The unique biological properties of carotenoids, which influence the immune system [6,8], intercellular communication control, differentiationcells growth regulation [9] and apoptosis [10] have activated the interest in this group of compounds for many years and are crucial for the understanding of their metabolism and beneficial role for human health [11].

Because of many antioxidative properties of carotenoids, the role of this class of compounds in modulation of the cancerogenesis process also focused interest of many researchers. Plant foods were shown to be inversely associated with cancer risk in epidemiologic studies [12–14]. The significant association with cancer risk has been reported for α-carotene, β-carotene and lycopene with breast cancer [15–22], and with ovarian cancer [23,24], for lycopene with prostate cancer [25,26], for lycopene, lutein, vitamin E and β-carotene with colorectal cancer [27–34]. However, not all reported results are consistent and more studies are needed to find clear relation between cancer risk and supplementation using carotenoids [35–37].

The purpose of this study was to show that the protective role of β-carotene against reactive oxygen species (ROS) can be investigated based on analysis of human colon single cells by using Raman spectroscopy and imaging. The influence of β-carotene supplementation time was also taken into cognizance.

The results for normal human colon cells, normal human colon cells exposed to oxidative stress conditions by using tBHP, normal human colon cells upon of β-carotene supplementation and tBHP adding and human colon cancer cells were compared.

## Materials and methods

### Cell culture

CCD-18Co (CRL-1459) cell line was purchased from ATCC and cultured using ATCC-formulated Eagle’s Minimum Essential Medium, Catalog No. 30-2003. To make the complete growth medium, the fetal bovine serum to a final concentration of 10% was added. Every 2 to 3 days the new medium was used. CaCo-2 cell line was also purchased from ATCC and cultured according to the ATCC protocols. The base medium for this cell line was ATCC-formulated Eagle’s Minimum Essential Medium, Catalog No. 30-2003. To make the complete growth medium, we have added fetal bovine serum to a final concentration of 20%. Medium was renewed 1 to 2 times a week. Cells used in the experiments were stored in an incubator providing environmental conditions at 37 °C, 5% CO_2_, 95% air. For all results presented in this manuscript we have recorded the Raman spectra and imaging for paraformaldehyde fixed cells. The procedure for fixed cells was as follows: cells were seeded onto CaF_2_ windows (25 × 1 mm) at a low density of 10^3^ cells/cm^3^. After 24 h or 48 h incubation on the CaF_2_ slides with β-carotene (c=10 µM) the tBHP for a final concentration of 200 µM was added for 30 min, after a half hour the cells were rinsed with phosphate-buffered saline (PBS,SIGMA P-5368, pH 7.4 at 25 °C, c= 0.01 M) to remove any residual medium or/and any residual β-carotene, fixed in paraformaldehyde (4% buffered formaldehyde) for 10 minutes and washed twice with distilled water. The Raman confocal measurements were performed immediately after the preparation of the samples.

### Raman spectra and imaging acquisition and analysis

All Raman images and spectra reported in this manuscript were recorded using the alpha 300 RSA+ confocal microscope (WITec, Ulm, Germany) using a 50 μm core diameter fiber, 532 nm excitation line, an imaging spectrograph/monochromator (Acton-SP-2300i), a CCD camera (Andor Newton DU970-UVB-353) and an Ultra High Throughput Spectrometer (UHTS 300). Excitation laser line was focused on the samples through a 40 x water dipping objective (NA: 1.0). The average laser excitation power was 10 mW, with integration time of 1.0 sec for low-frequency region and 0.5 sec for high-frequency region. Edge filters were used to remove the Rayleigh scattered light. A piezoelectric table was used to record Raman images. Spectra were collected at one acquisition per pixel. The cosmic rays were removed from each Raman spectrum (model: filter size: 2, dynamic factor: 10) and the Savitzky–Golay smoothing procedure was also implemented (model: order: 4, derivative: 0). Data acquisition and processing were performed using WITec Project Plus software.

All imaging data were analyzed using the Cluster Analysis method (CA) implemented in WITec Project Plus software. Briefly, CA allows to group objects. Vibrational spectra in our studies were analyzed in such a way that in the same group called a cluster were gathered all Raman spectra resembling each other, Raman spectra characterize by different vibrational features created another cluster. CA was performed using WITec Project Plus model (Centroid) with the k-means algorithm (each cluster was represented by a single mean vector). The normalization was performed using Origin software (model: divided by the norm).

### Chemical compounds

β-carotene catalogue Number C9750, Luperox^®^ TBH70X, tert-Butyl hydroperoxide solution catalogue Number 458139, bisBenzimide H 33342 trihydrochloride catalogue Number B2261, Red Oil-O catalogue Number O0625 were purchased from Sigma-Aldrich, and used without additional purification. XTT proliferation Kit with catalogue Number 20-300-1000 was purchased from Biological Industries.

### XTT cells viability tests

Tetrazolium salts have been widely used for many years as detection reagents in histochemical and cell biology tests [38,39]. The second generation tetrazolium dye, XTT, can be effectively used in tests to assess viability and proliferation, metabolic cytotoxicity, respiratory chain activity and apoptic cells [39–41].

The use of the XTT reagent is based on the statistical calculation of the metabolic activity of cells using a colorimetric technique which is a reaction to the changed environmental conditions in which they are located. The XTT test requires the use of a wavelength of 450 nm, from which the specific signal of the sample is obtained, and 650 nm constituting the reference sample during the test. The XTT test was used to calculate the metabolic activity of a living cell.

Tetrazolium salts are converted in cells with the help of special enzymes called doformazan, however, this reaction occurs properly only in cells with undamaged metabolism. The XTT test counts the final amount of formazan and converts it to the number of live cells. The advantage of the XTT test is the fact that this reagent is soluble in water, thanks to which the first representative readings can be obtained after about 3 hours of incubation. The sensitivity of the XTT test can be significantly increased thanks to the use of an intermediate electron carrier - PMS (n-methyldibenzopyrazine methyl sulfate) - which catalyzes the reduction of XTT and formation of a formazan derivative. The PMS activation reagent is included with the XTT in the ATCC XTT Cell Proliferation Assay Kit (ATCC^®^30-1011K ™) that was used for the cell culture studies in this work.

XTT tests on cell lines were carried out on a multi-sensing BioTek Synergy HT model reader designed for microplate testing. The test protocol was created especially for the presented experiments. After the microplate was applied to the reader, the inside of which was at a room temperature of 20-22°C, the sample underwent a pre-mixing process. Then, the actual measurement was carried out, consisting in examining the signal coming from the sample with excitation at 450 nm. After measuring the samples, the instrument carried out a reference measurement with excitation at 650 nm wavelength. After this sequence the microplate was removed from the reader.

The stress compound solutions were prepared by pre-diluting it in the PBS solvent. After obtaining the initial concentration of 20,000 μM, it was diluted in a medium intended for a given type of cells, so as to obtain the final concentration of 200 μM.

Scheme 1 shows the results of XTT test obtained for CCD-18Co human, normal, colon cells incubated with β-carotene in various concentrations and in various time.

**Scheme 1.**
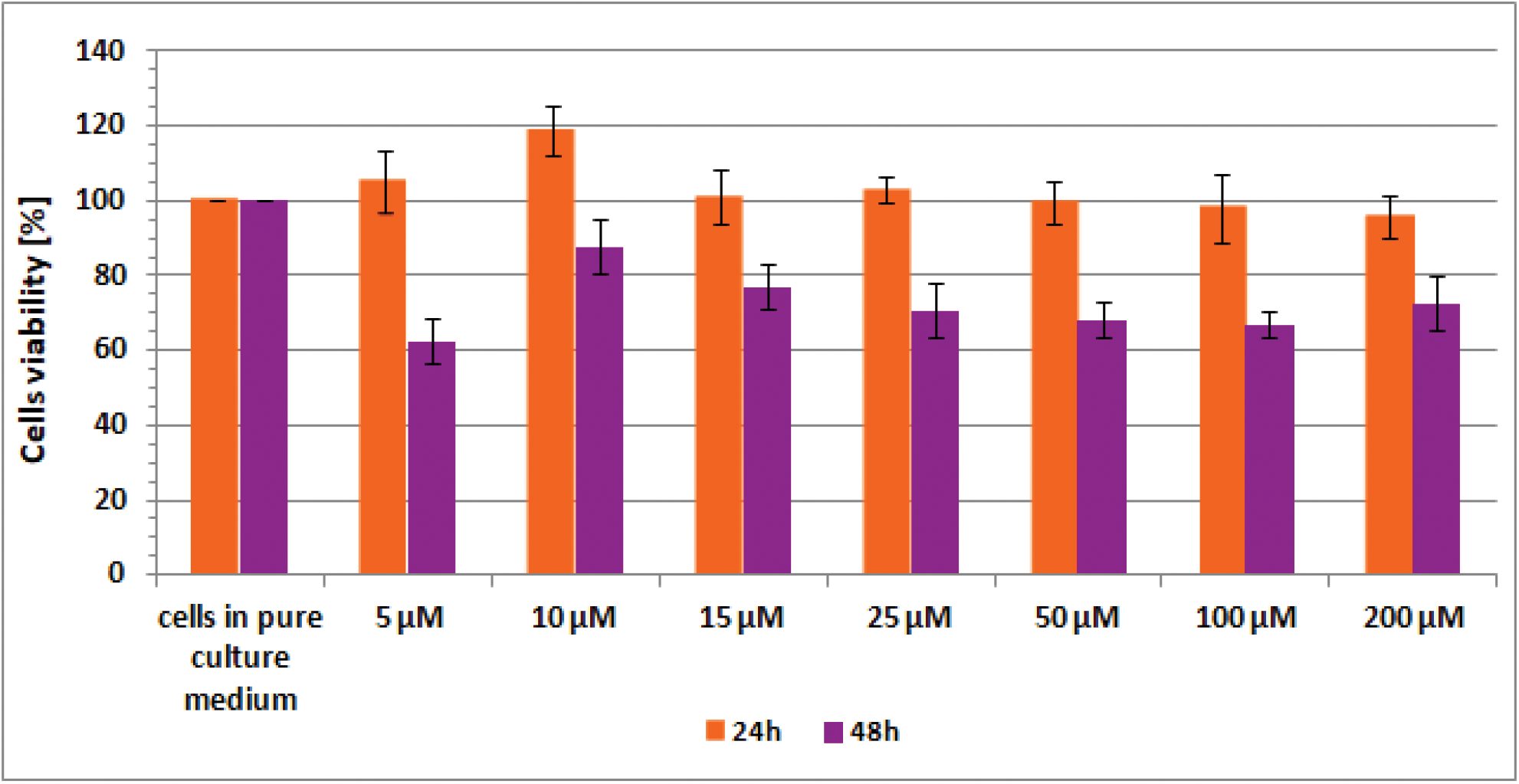
Results of XTT analysis obtained for CCD-18Co human, normal, colon cells supplemented with different concentrations of β-carotene in different time intervals.

## Results

In this section, the data obtained by using Raman spectroscopy and Raman imaging for human normal colon cells before and after ROS generation, including cells supplemented with β-carotene before ROS injuring are presented. We will show also the comparison on results obtained for normal human colon cells CCD-18Co in oxidative stress conditions and cancer human colon cell line CaCo-2.

Generally, in the Raman vibrational spectra there are two regions of interest: the Raman fingerprint region: 500–1800 cm^-1^ and the high frequency region: 2700–3100 cm^-1^ (the region 1800-2700 cm^-1^ is not considered for analysis due to the lack of Raman bands).

Figure 1 presents the microscopy image, Raman image of single human normal colon cell CCD-18Co constructed based on Cluster Analysis (CA) method, Raman images of all clusters identified by CA assigned to: nucleus, mitochondria, lipid-rich regions, membrane, cytoplasm, and cell environment, fluorescence images of lipid-rich regions and nucleus, the average Raman spectrum typical for single human normal colon cell CCD-18Co presented on microscopy image, the average Raman spectra typical for CCD-18Co human normal colon cells mean ± SD for all identified clusters for low frequency and high frequency region, and the average Raman spectrum for human normal colon cells mean ± SD for cells as a whole, all data for experiments performed without any supplementation of β-carotene, cells measured in PBS, colors of the spectra correspond to the colors of clusters.

**Figure 1.**
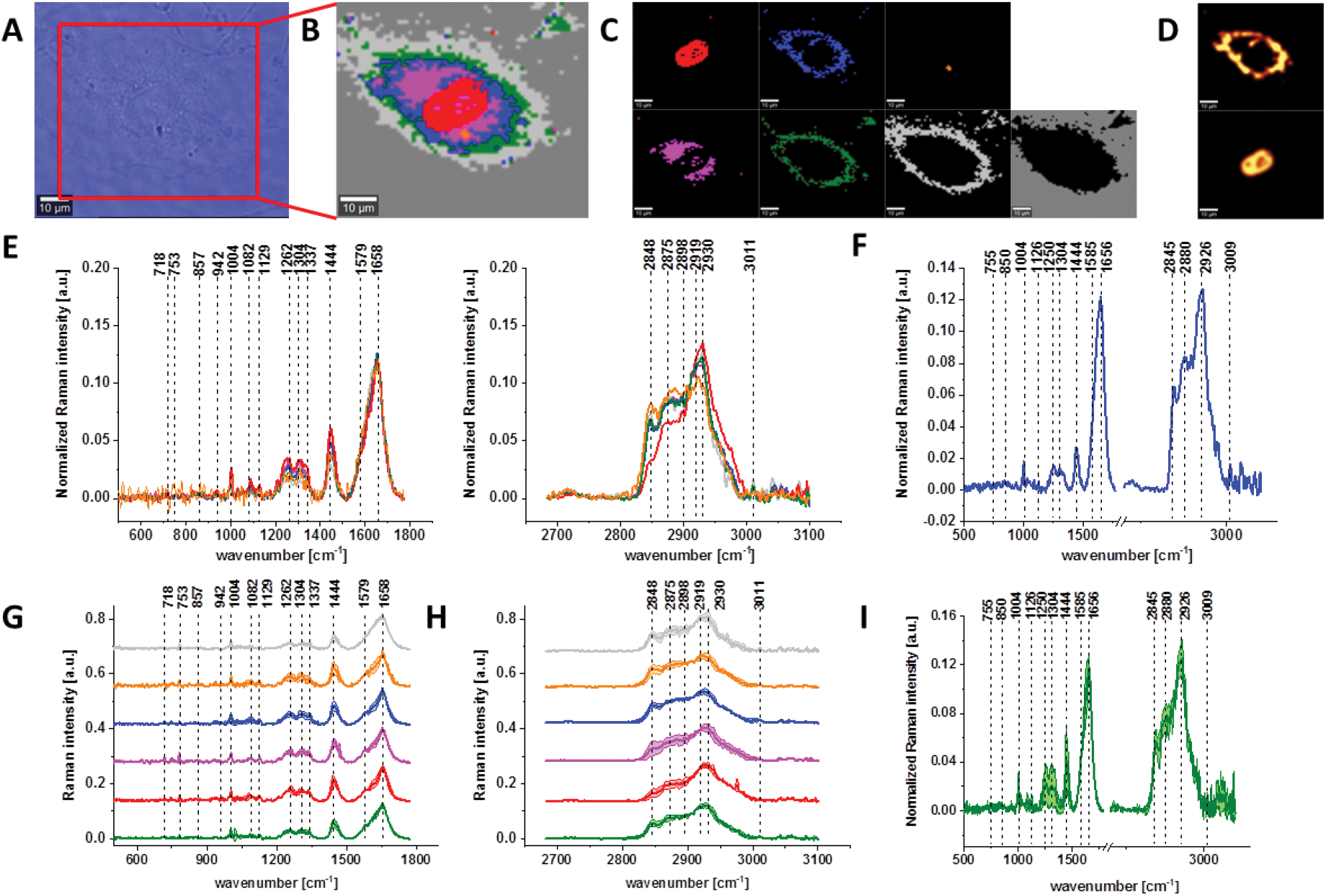
The microscopy image (A), Raman image of single human normal colon cell CCD-18Co constructed based on Cluster Analysis (CA) method (B), Raman images of all clusters identified by CA assigned to: nucleus (red), mitochondria (magenta), lipid-rich regions (blue, orange), membrane (light grey), cytoplasm (green), and cell environment (dark grey) (C), fluorescence images of lipid-rich regions (top panel) and nucleus (bottom panel) (D), the average Raman spectra typical for all clusters identified by CA for single human normal colon cell CCD-18Co (E), the average Raman spectrum typical for CCD-18Co single cell as a whole (F), the average Raman spectra typical for all clusters identified by CA typical for CCD-18-Co cells mean ± SD: nucleus (red), mitochondria (magenta), lipid-rich regions (blue, orange), membrane (light grey), cytoplasm (green), and cell environment (dark grey) for low-(G) and high-frequency region (H), the average Raman spectrum typical for CCD-18Co cells as a whole, mean ± SD (I), number of cells n=3, all data for experiments performed without any supplementation of β-carotene, cells measured in PBS, colors of the spectra correspond to the colors of clusters, excitation wavelength 532nm.

Because β-carotene is soluble only in organic compounds, we decided to compare the Raman spectra typical for CCD-18-Co human normal colon cells analyzed in PBS (shown in Figure 1) with results obtained for CCD-18-Co human normal colon cells after adding the same volume of organic solvent (THF/EtOH) as for 10 μM β-carotene supplementation (10 μM it was the final concentration obtained in culturing medium). The results of Raman imaging analysis of CCD-18Co human colon normal cells after adding the mixture of organic solvents to PBS including the comparison on the average Raman spectra typical for cells as whole analyzed in PBS and in PBS/THF/EtOH are shown on Figure 2.

**Figure 2.**
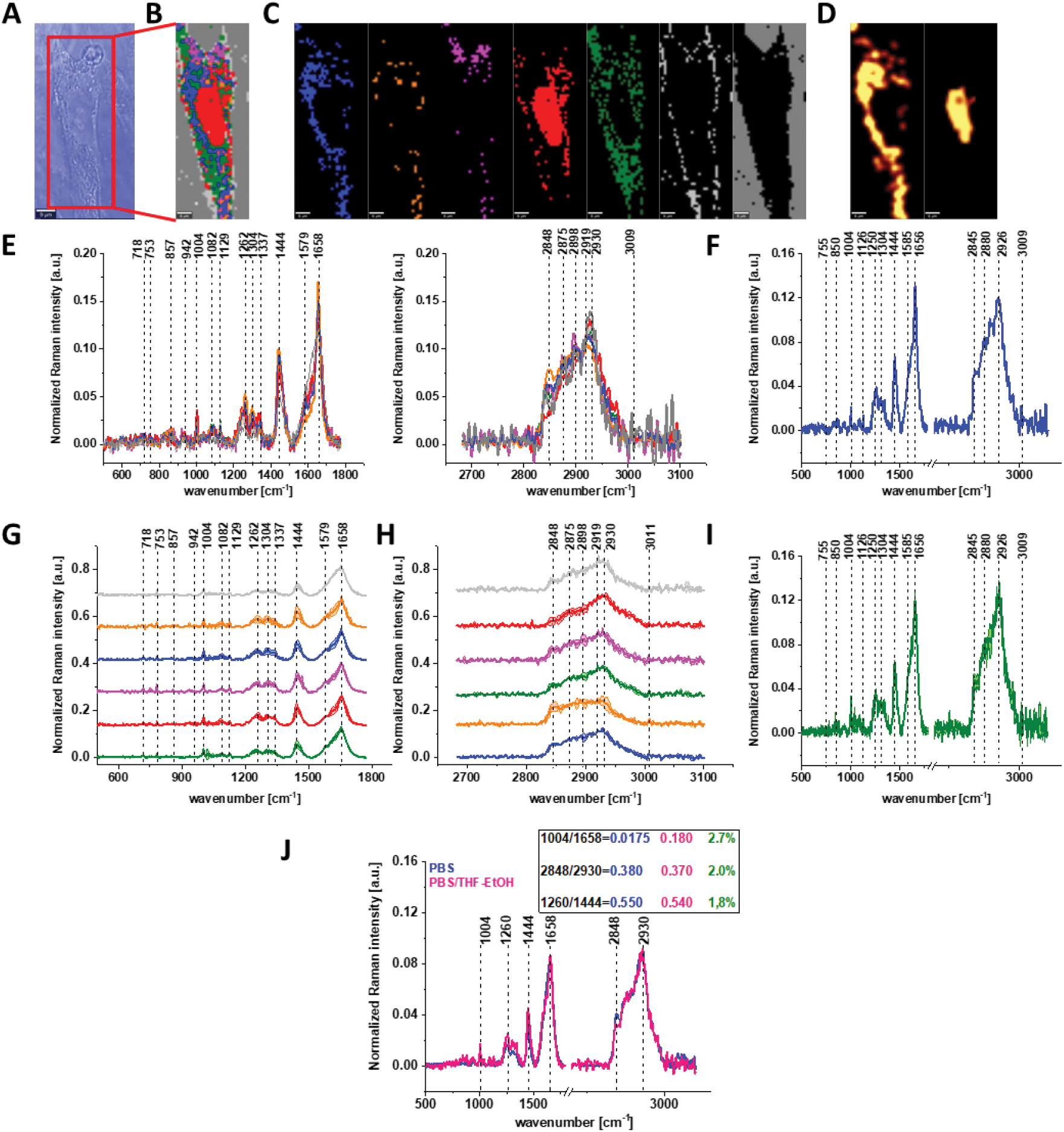
The microscopy image (A), Raman image constructed based on Cluster Analysis (CA) method (B), Raman images of all clusters identified by CA assigned to: nucleus (red), mitochondria (magenta), lipid-rich regions (blue, orange), membrane (light grey), cytoplasm (green), and cell environment (dark grey) (C), fluorescence images of lipid-rich regions (left panel) and nucleus (right panel) (D), the average Raman spectra typical of all clusters identified by CA for human colon normal single cell CCD-18Co (E), the average Raman spectrum typical for single human colon normal cell CCD-18Co as the whole (F), the average Raman spectra typical for all clusters identified by CA typical for CCD-18-Co cells mean ± SD: nucleus (red), mitochondria (magenta), lipid-rich regions (blue, orange), membrane (light grey), cytoplasm (green), and cell environment (dark grey) for low-(G) and high-frequency region (H), the average Raman spectrum typical for CCD-18Co cells mean ± SD (I), number of cells n=3, colors of the spectra correspond to the colors of clusters, all data for cells measured in PBS/THF/EtOH solution, excitation wavelength 532 nm, and the comparison of the average spectra for CCD-18-Co human normal colon cells in PBS/THF/EtOH – pink and in pure PBS – blue with calculated Raman bands ratios, the percentage difference between Raman peaks ratios based on average cells spectra is shown in green (J).

One can see from Figure 2 that the differences between the average Raman spectra (for cells as a whole) recorded in PBS and in PBS/THF/EtOH solution are subtle and fluctuate around 2%. Even though the difference is very slight, as the control group in all further comparisons and calculations we used results obtained for CCD-18-Co cells in PBS/THF/EtOH solution.

Results presented in Figures 1 and 2 confirm that Raman spectroscopy and imaging can be used to characterize the biochemical composition of human normal colon cells. Table 1 shows the main chemical components which can be identified based on their vibrational features in analyzed cells and the tendency in Raman peaks intensities observed for CCD-18Co human, normal colon cells in oxidative stress conditions generated by tBHP adding (discussed later in the manuscript).

**Table 1.** Band positions, tentative assignments for human normal colon cells from control sample (CCD-18Co in PBS/THF/ETOH) and the tendency in Raman peaks intensities observed for CCD-18Co human, normal colon cells in oxidative stress conditions generated by tBHP adding (discussed later in the manuscript). Data based on the average Raman spectra for cells as a whole [42].

| Wavenumber<br>[cm<sup>-1</sup>] | Tentative assignments of Raman bands | Tendency in Raman peak<br>intensity observed for CCD-18Co human, normal colon cells<br>in oxidative stress conditions<br>generated by adding tBHP |
| --- | --- | --- |
| 716 | C-N (membrane phospholipids head)/adenine ,<br>CN<sub>2</sub>(CH<sub>3</sub>)<sub>3</sub> (lipids), choline group | ↘ |
| 812 | Tyrosine | ↘ |
| 820 | Structural protein modes of tumors, proteins,<br>including collagen I | ↘ |
| 832 | Tyrosine (Fermi resonance of ring fundamental<br>and overtone), asymmetric O-P-O stretching,<br>tyrosine | ↘ |
| 869 | C-C stretching modes of protein | ↘ |
| 891 | Protein bands, structural protein modes of tumors | ↘ |
| 932 | Skeletal C-C, α -helix, proline, hydroxyproline,<br>ν(C-C) skeletal of | ↘ |
|  | collagen backbon |  |
| **1004** | Symmetric ring breathing of phenylalanine | ↘ |
| **1078** | Symmetric PO<sub>2</sub><sup>-</sup> of DNA (represents more DNA in cell) | ↘ |
| **1254** | Amide III, adenine, cytosine | ↘ |
| **1304** | CH<sub>2</sub> deformation (lipid), adenine, cytosine | ↘ |
| **1444** | δ(CH<sub>2</sub>), lipids, fatty acids | ↘ |
| **1602** | δ(C=C), phenylalanine (protein assignment) | ↘ |
| **1626** | Amide C=O stretching absorption for the b-form polypeptide films | ↘ |
| **1654** | Amide I | ↘ |
| **1720** | C=O | ↘ |
| **1754** | C=O (lipid) | ↘ |
| **2854** | CH<sub>2</sub> symmetric stretch of lipids & CH<sub>2</sub> asymmetric stretch of lipids and proteins | ↘ |
| **2880** | CH<sub>2</sub> asymmetric stretch of lipids and proteins | ↘ |
| **2926** | Symmetric CH<sub>3</sub> stretch, due primarily to protein | ↘ |
| **3009** | v<sub>as</sub>(=C-H), lipids, fatty acids | ↘ |

The various signals seen on Figures 1 and 2 have been assigned to nucleic acids, proteins or lipids and provide adequate information to assess variations in spectral characteristics.

As we mentioned above the main goal of our experiments was to prove the protective role of natural antioxidants in oxidative stress conditions by using Raman spectroscopy and imaging, therefore the next step was the analysis of human colon normal cells CCD-18-Co after ROS generation. The main advantage of Raman imaging method that should be highlighted in this point is that vibrational imaging allows to investigate biochemical composition of cells without any cells destruction. Moreover, the high spatial resolution allows to track any changes on subcellular level.

Before we start the analysis of spectroscopic data we have to underline a few regularities.

ROS are produced by living organisms as a result of natural cellular metabolism, but in such a case the concentrations of them is low to moderate their functions in physiological cell processes, but at high ROS concentrations adverse modifications to cell components, such as: lipids, proteins, and DNA can be noticed [43–48]. The shift in balance between oxidant/antioxidant in favor of oxidants seems to be crucial for understanding many dysfunctions of human cells.

It has been shown in literature that oxidative stress contributes to many pathological conditions, including cancer [14–36], neurological disorders [49–52], atherosclerosis, hypertension, ischemia/perfusion [53–56], acute respiratory distress syndrome, idiopathic pulmonary fibrosis, chronic obstructive pulmonary disease [57], and asthma [58–60].

Generally, ROS can be divided into two groups: free radicals and nonradicals. Free radicals contain one or more unpaired electrons and are characterize by high reactivity to molecules. Free radicals can recombine sharing electrons and create nonradical forms. The three ROS of physiological significance are: superoxide anion, hydroxyl radical, and hydrogen peroxide. Superoxide anion is formed by the addition of one electron to the molecular oxygen. This process is mediated by nicotine adenine dinucleotide phosphate [NAD(P)H] oxidase, xanthine oxidase or by mitochondrial electron transport system. The main place where the superoxide anion is produced are mitochondria because 1-3% of all electrons “leak” from respiration chain system and produce superoxide. Superoxide is then converted into hydrogen peroxide, which diffuses across the plasma membrane by the action of superoxide dismutases.

The most reactive of ROS is hydroxyl radical, which can damage proteins, lipids, carbohydrates and DNA. Hydroxyl radical can start lipid peroxidation by taking an electron from polyunsaturated fatty acids. Other oxygen-derived free radicals are the peroxylradicals (ROO^·^(. Simplest form of these radicals is hydroperoxyl radical (HOO^·^), which plays a crucial role in fatty acid peroxidation. Free radicals can trigger lipid peroxidation chain reactions by abstracting a hydrogen atom from a sidechain methylene carbon. The lipid radical then reacts with oxygen to produce peroxyl radical. Peroxyl radical initiates a chain reaction and transforms polyunsaturated fatty acids into lipid hydroperoxides [61]. Lipid hydroperoxides are very unstable and easily decompose to secondary products, such as aldehydes. Isoprostanes are another group of lipid peroxidation products that are generated via the peroxidation of unsaturated fatty acids e.g. arachidonic acid.

Tert-Butyl hydroperoxide (tBHP) used in our experiments to produce ROS is well-known as a model substance for oxidative stress generation and subsequent analysis of cellular alterations in cells and tissues. Generally, in living organisms two pathways by which tBHP is metabolized can be distinguish; both of them induce oxidative stress. The first one provided by cytochrome P450, leads to production of peroxyl and alkoxyl radicals [62]. The second pathway employs glutathione peroxidase. Tert-Butyl hydroperoxide is detoxified to tert-butanol and reduced glutathione is depleted by oxidation to its disulphide form [21].

Figure 3 shows the microscopic image, Raman image constructed based on Cluster Analysis (CA) method, Raman images of all clusters identified by CA assigned to: nucleus, mitochondria, lipid-rich regions, membrane, cytoplasm, and cell environment, fluorescence images of lipid-rich regions and nucleus, the average Raman spectra typical of all clusters identified by CA for single human normal colon cell for low frequency and high frequency region, the average Raman spectrum typical of single CCD-18Co cell as a whole, and the average Raman spectra for human normal colon CCD-18Co cells mean ± SD after adding tBHP for 30 minutes, colors of the spectra correspond to the colors of clusters.

**Figure 3.**
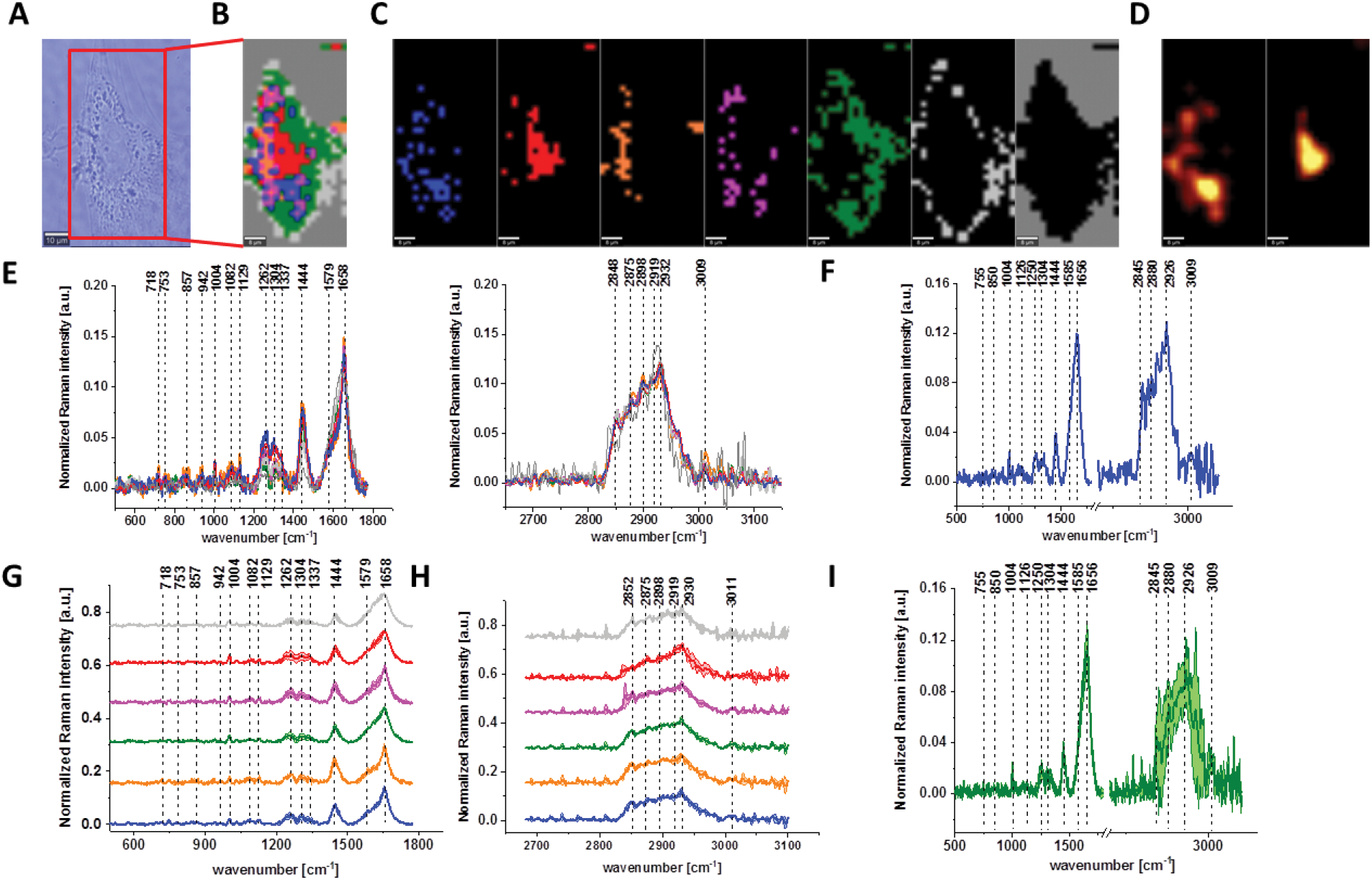
The microscopy image (A), Raman image constructed based on Cluster Analysis (CA) method (B), Raman images of all clusters identified by CA assigned to: nucleus (red), mitochondria (magenta), lipid-rich regions (blue, orange), membrane (light grey), cytoplasm (green), and cell environment (dark grey) (C), fluorescence images of lipid-rich regions (left panel) and nucleus (right panel) (D), average Raman spectra typical for all clusters identified by CA for single human normal colon cell (E), average Raman spectrum typical for CCD-18Co single cell as a whole (F), average Raman spectra typical for all clusters identified by CA typical for CCD-18-Co cells mean ± SD: nucleus (red), mitochondria (magenta), lipid-rich regions (blue, orange), membrane (light grey), cytoplasm (green), and cell environment (dark grey) for low-(G) and high-frequency region (H), average Raman spectrum typical for CCD- 18Co cells as a whole mean ± SD (I) after adding of tBHP for 30 minutes (final concentration in medium c=200 μM), number of cells n=3, colors of the spectra correspond to the colors of clusters, all data performed for PBS/THF/EtOH solution, excitation wavelength 532 nm.

Based on high resolved Raman spectra recorded for normal human colon cells CCD-18Co and for individual organelles without and after adding tBHP for ROS generation shown in Figures 2 and 3 one can obtained the complex information about the biochemistry of single cell as a whole or we can track changes in cell’s nucleus, mitochondria, and lipid structures. Figure 4 and Table 1 show the main spectroscopic differences detected based on vibrational Raman spectra before and after oxidative stress generation.

**Figure 4.**
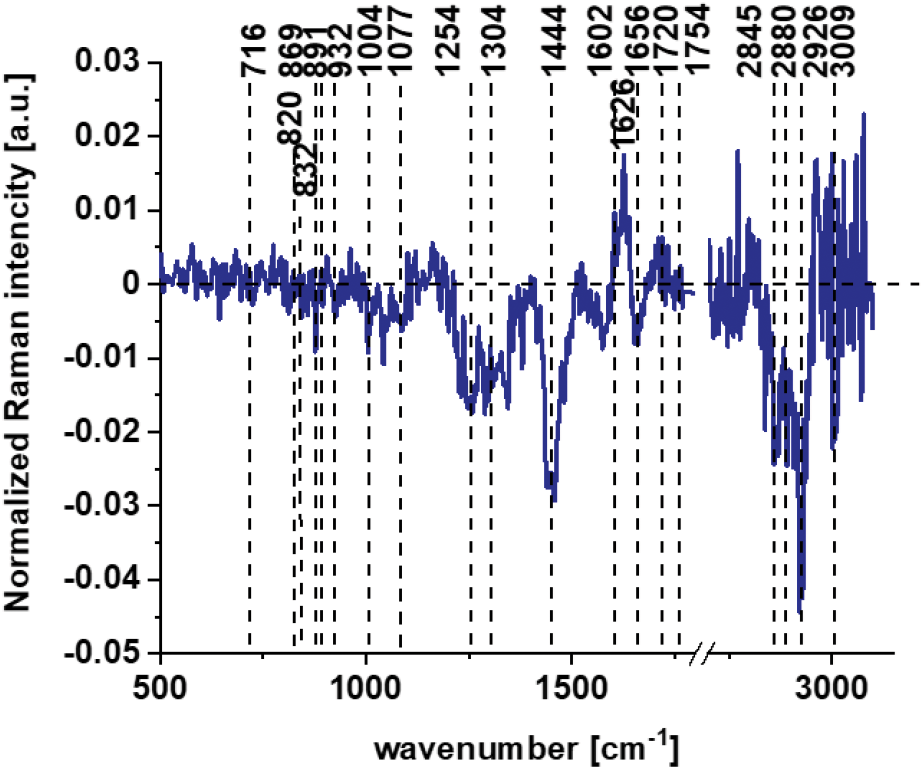
The difference spectrum calculated for human colon normal cells CCD18-Co control group and human normal colon cells CCD18-Co in oxidative stress conditions generated by tBHP adding (data based on the Raman vibrational spectra for cells as a whole).

Figures 4 shows the difference Raman spectrum of human normal colon cells from control sample (CCD-18Co in PBS/THF/ETOH solution) and human normal colon cells in oxidative stress conditions generated by tBHP adding (data based on the Raman vibrational spectra for cells as a whole).

One can see from Figure 4 that ROS generation has changed the biochemical composition of normal human colon cells CCD18-Co. In detail changes noticed in Figure 4 will be discussed in the manuscript later. The tendency in Raman peaks intensities observed for CCD-18Co human, normal colon cells after ROS generation by tBHP adding for cells as a whole is also shown in Table 1. As we have stressed above Raman microspecroscopy and Cluster Analysis of Raman data allow to observed the influence of ROS generation on individual organelles of human, normal, CCD18-Co cells. Figure 5 shows the difference spectra (green) calculated for nucleus, mitochondria, lipids structures, cell membrane and cytoplasm in high frequency region (A) and in low frequency region (B) based on the average Raman spectra typical for all clusters assigned to the subcellular structures for cells in control group (blue) and after tBHP adding (red).

**Figure 5.**
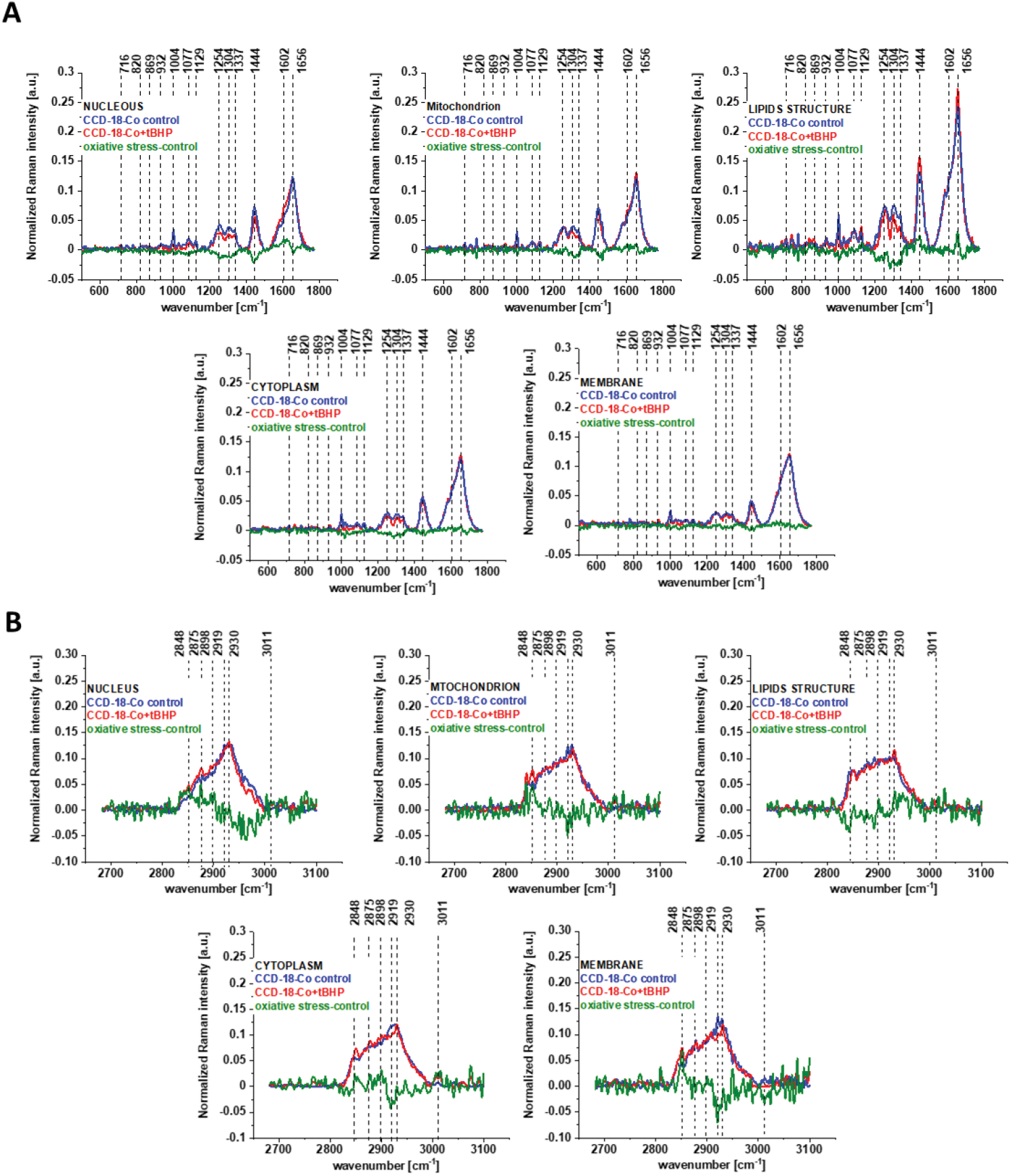
The difference spectra (green) calculated for nucleus, mitochondria, lipids structures, cell membrane and cytoplasm in low frequency region (A) and in high frequency region (B) based on the average Raman spectra typical of clusters assigned to the subcellular structures for CCD-18-Co cells in control group (blue) and after adding tBHP (red).

Figure 5 shows that the generation of ROS affects the intensity of the Raman signal assigned to all selected organelles of human normal CCD18-Co colon cells. This observation confirms that ROS generation can affect the function of all cell substructures.

The next step of the analysis was the interpretation of Ramana data obtained for CCD18-Co human normal colon cells at first supplemented by β-carotene for 24 or 48 hours and then treated using tBHP for 30 min.

Figure 6 shows all results obtained for human normal colon cells CCD-18Co after 24 h of β - carotene supplementation and then adding of tBHP for 30 min.

**Figure 6.**
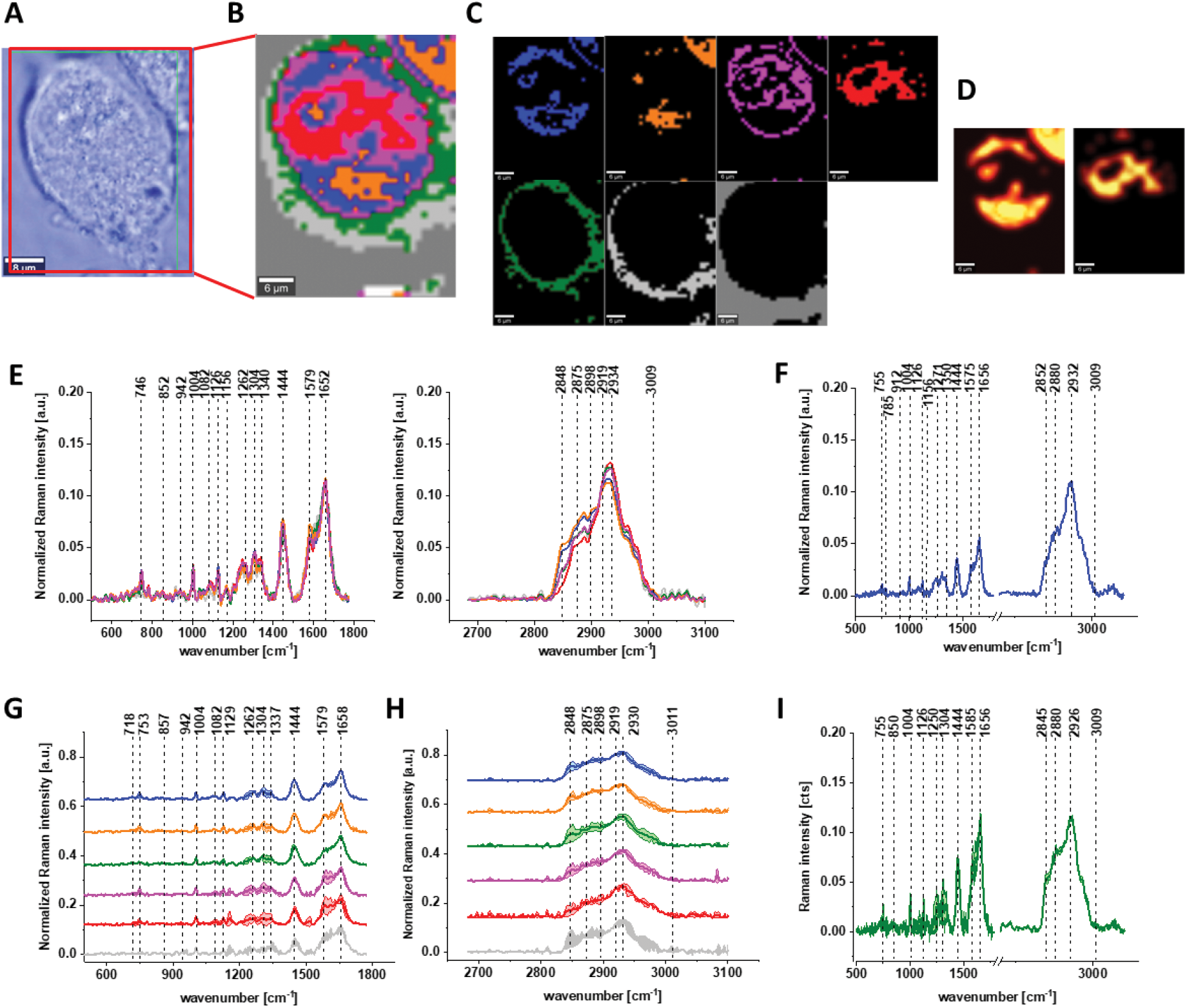
The microscopy image (A), Raman image constructed based on Cluster Analysis (CA) method (B), Raman images of all clusters identified by CA assigned to: nucleus (red), mitochondria (magenta), lipid-rich regions (blue, orange), membrane (light grey), cytoplasm (green), and cell environment (dark grey) (C), fluorescence images of lipid-rich regions (left panel) and nucleus (right panel) (D), average Raman spectra typical for all clusters identified by CA for single human colon cell (E), average Raman spectrum typical for CCD-18Co single cell as a whole (F), average Raman spectra typical for all clusters identified by CA typical for CCD-18-Co cells mean ± SD: nucleus (red), mitochondria (magenta), lipid-rich regions (blue, orange), membrane (light grey), cytoplasm (green), and cell environment (dark grey) for low-G) and high-frequency region (H), average Raman spectrum typical for CCD-18Co cells as a whole mean ± SD (I) for human normal colon cells CCD-18Co after at first β -carotene supplementation for 24 hours and then adding of tBHP for 30 minutes, number of cells n=3, colors of the spectra correspond to the colors of clusters, all data obtained for PBS/THF/EtOH solution, excitation wavelength 532 nm.

Figure 7 shows all results obtained for human normal colon cells CCD-18Co after 48 h of β- carotene supplementation and then adding of tBHP for 30 min.

**Figure 7.**
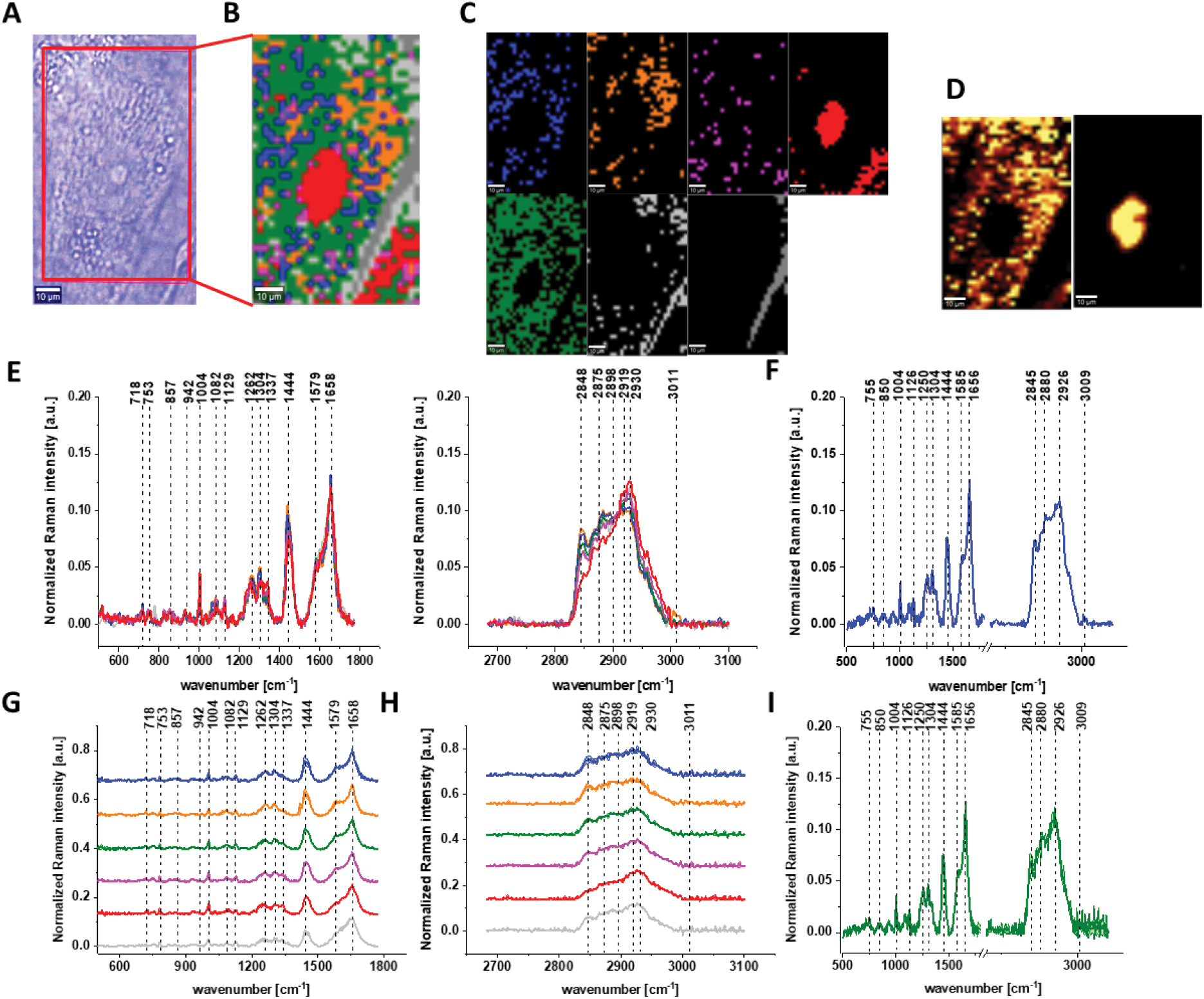
The microscopy image (A), Raman image constructed based on Cluster Analysis (CA) method (B), Raman images of all clusters identified by CA assigned to: nucleus (red), mitochondria (magenta), lipid-rich regions (blue, orange), membrane (light grey), cytoplasm (green), and cell environment (dark grey) (C), fluorescence images of lipid-rich regions (left panel) and nucleus (right panel) (D), average Raman spectra typical for all clusters identified by CA for single human colon cell (E), average Raman spectrum typical for CCD-18Co single cell as a whole (F), average Raman spectra typical for all clusters identified by CA typical for CCD-18-Co cells mean ± SD: nucleus (red), mitochondria (magenta), lipid-rich regions (blue, orange), membrane (light grey), cytoplasm (green), and cell environment (dark grey) for low-(G) and high-frequency region (H), average Raman spectrum typical for CCD-18Co cells as a whole mean ± SD (I) for human normal colon cells CCD-18Co after at first β-carotene supplementation for 48 hours and then adding of tBHP for 30 minutes, number of cells n=3, colors of the spectra correspond to the colors of clusters, all data for PBS/THF/EtOH solution.

One can see form Figures 6 and 7 that also for conditions including β-carotene supplementation and ROS generation we can obtain high resolved Raman spectra based on which the influence on biochemistry of single cells of antioxidant supplementation and oxidative stress can be analyzed.

## Discussion

Having reached this point when the analysis based on Raman vibrational spectra for different human normal colon CCD-18Co cells groups has been performed we can compare and discuse results shown on Figures 1-7.

Considering that CCD18-Co human normal colon cells are basically composed of three types of macromolecules: proteins, nucleic acids and lipids, in order to explore characteristic changes under ROS conditions we will focused the attention to these class of compounds. The qualitative and quantitative comparison between paired bands assigned to proteins, lipids and nucleic acids will continue to be discussed according to the biological attribution of them.

Based on the results shown in Table 1 and frequencies highlighted in Figures 1-7 we have chosen the following Raman bands to compare: 812, 832, 869, 932, 1004, 1254, 1654 cm^-1^.

Figure 8 shows the comparison of Raman band intensity ratios for four different human normal colon cells groups: control group, group treated by using tBHP for 30 min, group at first supplemented with β-carotene for 24h and after that incubated with tBHP for 30 min, and group at first supplemented with β-carotene for 48 h and after that incubated with tBHP for 30 min.

**Figure 8.**
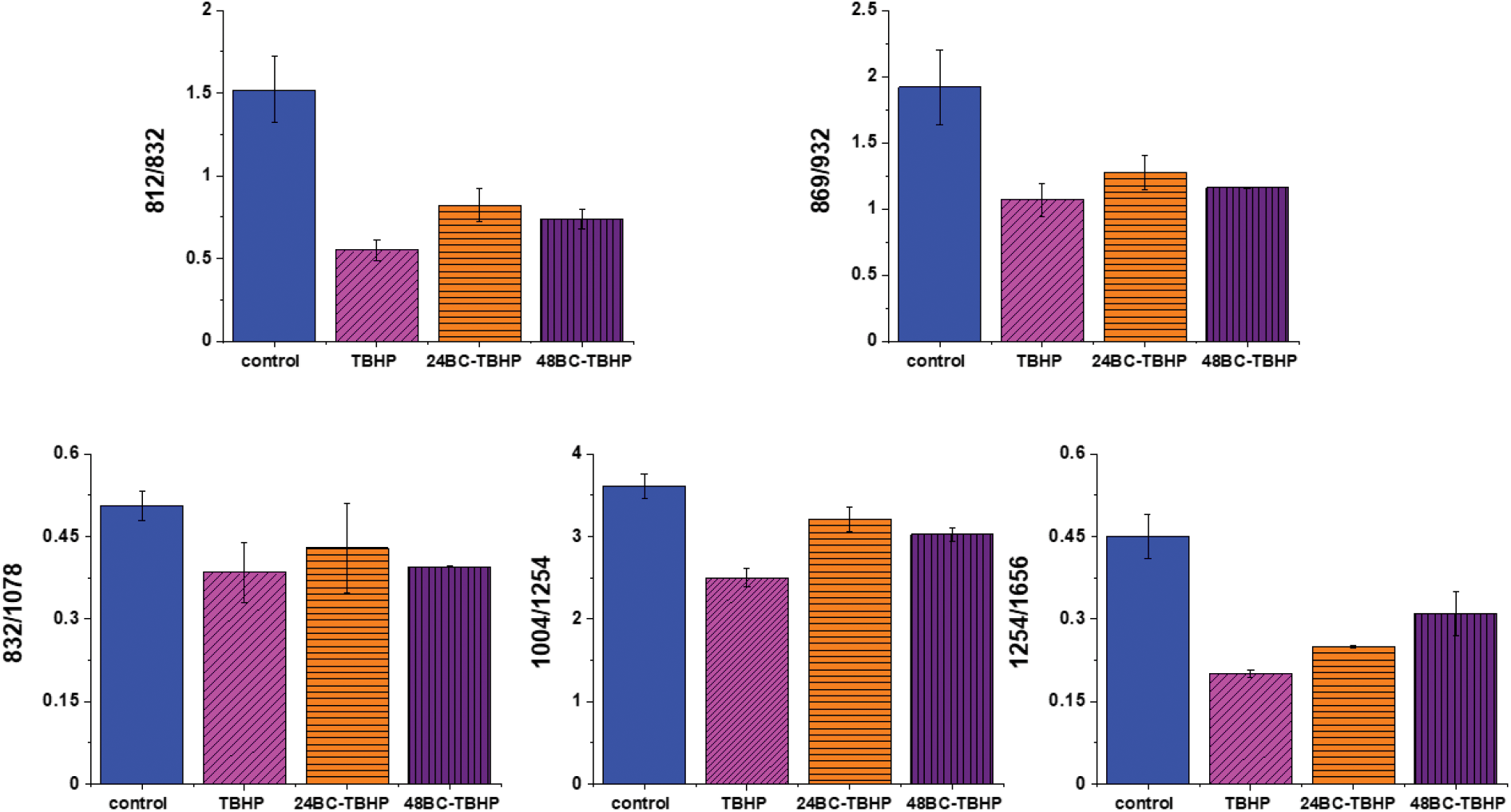
Raman band intensities ratios for selected Raman bands corresponding to proteins 812/832, 869/932, 832/1078, 1004/1254, 1254/1656 for four groups of normal human colon cells CCD18-Co: control group (labeled control, blue), group after ROS generation by using tBHP (labeled with TBHP, magenta), cells group at first supplemented with β-carotene for 24 h and then treated by tBHP (labeled 24BC-TBHP, orange), group at first supplemented with β- carotene for 48 h and then treated by tBHP (labeled 24BC-TBHP, violet).

Compared with the oxidation group (magenta), the relative intensities of several protein bands 812/832, 869/932, 1004/1254 and 832/1078 (the ratio for Raman bands typical for protein (tyrosine) and DNA) in the control (blue) and protective groups (at first supplemented by β-carotene, orange and violet) show the same variation trend.

In detail for Raman bands at 812 and 832 cm^-1^ for tBHP group (after ROS generation) compare to control one the decreasing for Raman bands ratio I_812_/I_832_ was observed. Decreasing observed in the oxidation group, indicates that tyrosine and its internal ‘out of plane ring breathing mode’ was affected by oxidation. Such interaction suggests that the molecular vibration of tyrosine is sensitive to oxidation, resulting in structural changes. The adding of β-carotene before ROS generation results in increasing of analyzed ratio compared to the oxidation group.

The bands at 869 and 932 cm^−1^ also correspond to proteins. The band 869 cm^-1^ came from C-C stretching modes of proteins, while the band at 932 cm^−1^ correspond to C-C stretching vibrational mode of proline, valine and protein backbone (α-helix conformation). One can see form Figure 8 that the ratio of Rama band intensities I_869_/I_932_ is significantly decreased in the oxidation group, and once again, increased after adding β-carotene. Although the observed increasing confirms the protective role of β-carotene, still the relative intensity is lower than observed for the control group. Summarizing, all observations made for the ratio I_869_/I_932_ indicate that tBHP changes the content and structure of some amino acids by reaction of ROS with proteins, resulting in concertation modulation and functional damage of them. This observation is consistent with data published in the literature [63]. The same trend was observed for the ratio calculated based on the Raman bands corresponding to phenylalanine (1004 cm^-1^) and Amide III. Once again one can observe the decreasing of the ratio for oxidation group and protective role of the antioxidant even if the values for 24BC+tBHP and 48BC+tBHP groups are lower than typical for control cells. The same variation for all analyzed cells groups was noticed for Raman bands 1245 and 1656 cm^-1^. The decreasing of the ratio I_1254_/I_1656_ for ROS injured cells and systematic increase of this ratio as a function of incubation time with β-carotene was observed. Moreover, the change of band ratio of amides I_1254_/I_1656_ may suggest also that a desamidization was triggered and resulted in modification of the spatial structure of proteins, in loss of protein activity or modification of their biological function [64].

The analysis of Raman bands ratios has been performed also for bands typical for nucleic acids and lipids. Figure 9 shows the results obtained for these types of compounds.

**Figure 9.**
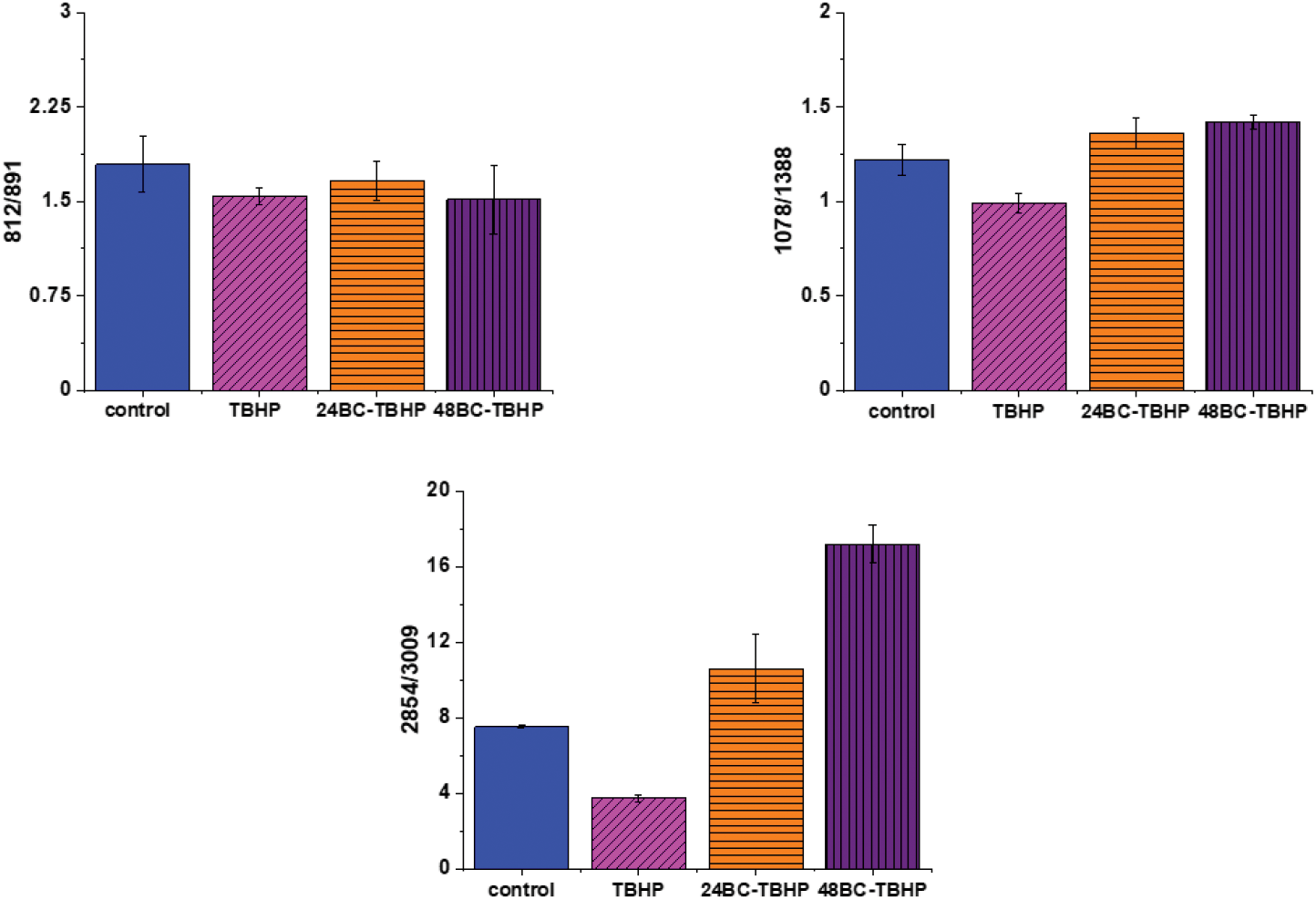
Raman band intensities ratios for selected Raman bands corresponding to nucleic acids 812/891, 1078/1368 and lipids 2854/3009 for four groups of normal human colon cells CCD18-Co: control group (labeled control, blue), group after ROS generation by using tBHP (labeled with TBHP, magenta), cells group at first supplemented with β-carotene for 24 h and then treated by tBHP (labeled 24BC-TBHP, orange), group at first supplemented with β- carotene for 48 h and then treated by tBHP (labeled 48BC-TBHP, violet).

For bands typical for nucleic acids (812, 891, 1078, 1388 cm^-1^) and lipids (2854, 3009 cm^-1^) once again as for bands typical for proteins we have noticed changes induced by ROS and modulations triggered by β-carotene supplementation. Lipids are the main component of subcellular structures like lipid droplets and the cell membrane, which is susceptible to peroxidation induced by ROS, Products from ROS reaction can cause damage to either the cell membrane or the organelle membrane [65,66]. The main protective role of β-carotene in this case was seen for lipids fraction, especially unsaturated one observed in high-frequency region. We have to remember that lipids peroxidation is a process that consists of three phases: initiation, propagation and termination. In the initiation phase, the hydrogen atom is separated from the molecule of polyunsaturated fatty acid or the rest of such acid that is part of the phospholipid under the influence of e.g. the hydroxyl radical In the propagation phase reactions alkyl free radicals react with oxygen to form peroxide free radicals, which detach hydrogen atoms from subsequent, undamaged molecules of unsaturated fatty acids. This reaction produces fatty acid peroxide and another alkyl radical which can be oxidized another molecule of fatty acid. The above reactions can repeat many times, which leads to the transformation into peroxides - several, several dozen or even several hundred fatty acid molecules. Lipid peroxidation products change the physical properties of all cell membranes. For phospholipids located inside the lipid bilayer, introduced polar peroxide, ketone and aldehyde groups remain or hydroxy. This lowers the hydrophobicity of the lipid interior of cell membranes, as well as changes the organization of the lipid bilayer, which to disturbance of lipid asymmetry of membranes [67].

We also noticed also that the β-carotene may affect the ROS damages induced in nucleic acids (812/893, 1078/1386)). DNA ROS damage effect may involve two aspects, nucleic acid bases and DNA phosphoric acid skeleton [68]. ROS are well known as mediators of double strand breaks (DSBs), which can be mutagenic due to chromosomal rearrangements or loss of genetic information due to erroneous DNA repair. ROS have also been reported to directly induce other forms of DNA damage through oxidizing nucleoside bases e.g. formation of 8-oxo guanine) [69], which can lead to G-T or G-A transversions if unrepaired. Oxidized bases are typically recognized and repaired by the Base Excision Repair (BER) pathway, but when they occur simultaneously on opposing strands, attempted BER can lead to the generation of DSBs [70]. It has been shown that ROS accumulation also induces mitochondrial DNA lesions, strand breaks and degradation of mitochondrial DNA [71]. Data presented on Figure 9 confirm that based on Raman spectra the changes in DNA after ROS generation can be tracked.

It has been shown that permanent changes in the genome of cells e.g. as a result of oxidative stress, it is the first stage characteristic of the process of mutagenesis, carcinogenesis and cell aging. The appearance of a mutation in DNA is a critical step in the process of cancer development and more than 100 DNA modifications have been identified in cancer cells products [72].

Finally, we have decided to compare the results obtained for CCD18-Co human normal colon cells in oxidative stress conditions with data obtained for CaCo-2 human cancer cell line. Figure 10 shows the data obtained for human cancer colon cells CaCo-2. Figure 11 the comparison for selected Raman bands typical for proteins, nucleic acids and lipids.

**Figure 10.**
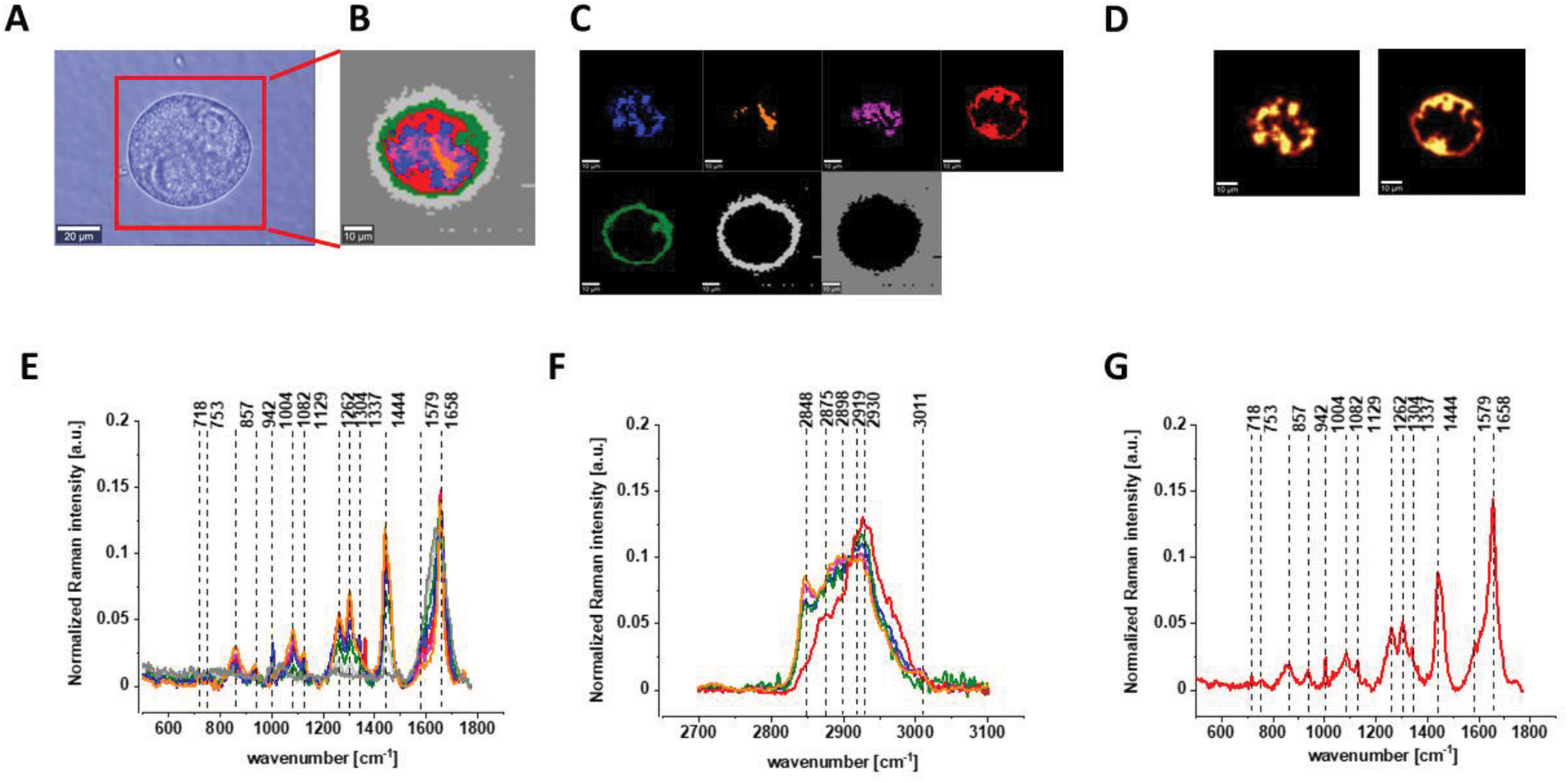
The microscopy image (A), Raman image constructed based on Cluster Analysis (CA) method (B), Raman images of all clusters identified by CA assigned to: nucleus (red), mitochondria (magenta), lipid-rich regions (blue, orange), membrane (light grey), cytoplasm (green), and cell environment (dark grey) (C), fluorescence images of lipid-rich regions (left panel) and nucleus (right panel) (D), average Raman spectra typical for all clusters identified by CA for single cancer human colon cell CaCo-2 in a low- (E) and high-frequency region (F), average Raman spectrum typical for CaCo-2 single cell as a whole (G), colors of the spectra correspond to the colors of clusters, all data for PBS/THF/EtOH solution.

**Figure 11.**
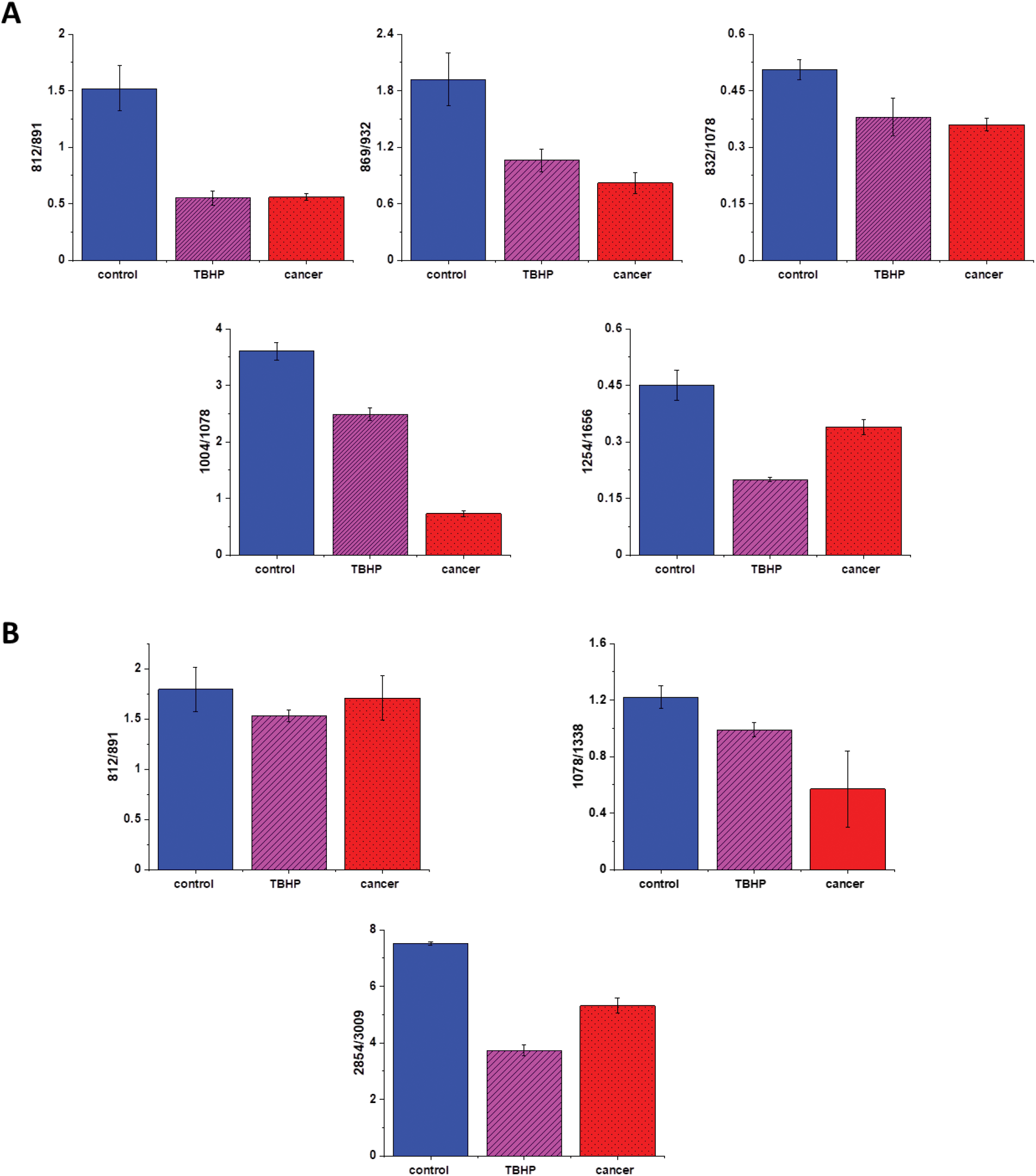
Raman band intensities ratios for selected Raman bands corresponding to proteins 812/832, 869/932, 832/1078, 1004/1254, 1254/1656, (A), nucleic acids lipids 812/891, 1078/1368 and 2854/3009 (B) for three groups of human colon cells: CCD18-Co cells - control group (labeled with control, blue), CCD18-Co after ROS generation by using tBHP (labeled with TBHP, magenta), and CaCo-2 human cancer cells (labeled CaCo-2, red).

Figure 11 shows the comparison of ratios for selected Raman bands typical for proteins, nucleic acids and lipids for CCD-18Co human normal colon cells, CCD-18Co human normal colon cells after ROS generation by adding of tBHP for 30 min. and CaCo-2 human cancer colon cells.

One can see form Figure 11 that intensities of Raman band typical for proteins, nucleic acids and lipids are comparable for CCD18-Co human normal colon cells in oxidative stress conditions and for CaCo-2 human colon cancer cell and simultaneously differ significantly from control group. This observation confirm that Raman band intensities analysis allow to track biochemical changes induced by ROS and during cancerogenesis. Moreover, comparable ratios for ROS injured cells and cancer cells may confirm that cells altered by ROS shows many common characteristic with pathological cells.

## Conclusions

In the presented work, we performed label-free detection of tBHP-induced oxidative stress by using Raman imaging and spectroscopy. We have shown that Raman imaging and spectroscopy are capable to characterize and differentiate human normal colon cells CCD-18Co, human normal colon cells after ROS generation, human normal colon cells at first incubated with β-carotene and after that treated by using tBHP, and finally human, cancer colon cells CaCo-2.

Moreover, we have confirmed that substructures of human colon single cells such as: nucleus, mitochondria lipid-rich regions, membrane, and cytoplasm can be precisely visualized based on Raman spectra. Moreover, biochemical changes typical for selected cells substructures can be tracked based on vibrational features.

We have shown also that fluorescence based images accurately correspond to the Raman images confirming localization of cells substructures such as: nucleus and lipids-rich structures.

Based on Raman band intensities attributed to proteins, nucleus acids and lipids as well as ratios 812/832, 869/932, 832/1078, 1004/1254, 1254/1656, 812/891, 1078/1368 2854/3009 calculated based on them, we have confirmed the protective role of β-carotene for cells in oxidative stress conditions for label-free and nondestructive spectroscopic method.

## Author Contribution

Conceptualization: BB-P; Funding acquisition: BB-P; Investigation: BB-P, KB; Methodology: BB-P, KB, Writing – original draft: BB-P; Manuscript editing: BB-P, KB All authors reviewed and provide feedback on the manuscripts.

